# Metaproteomics Reveals Potential Mechanisms by which Dietary Resistant Starch Supplementation Attenuates Chronic Kidney Disease Progression in Rats

**DOI:** 10.1101/340513

**Authors:** Boris L Zybailov, Galina V Glazko, Yasir Rahmatallah, Dmitri S Andreyev, Taylor McElroy, Oleg Karaduta, Stephanie D Byrum, Lisa Orr, Alan J Tackett, Samuel G Mackintosh, Ricky D Edmondson, Dorothy A Kieffer, R J Martin, Sean H Adams, Nicolas D Vaziri, John M Arthur

**Affiliations:** Department of Biomedical Informatics, UAMS, Little Rock, AR; Department of Biochemistry and Molecular Biology, UAMS, Little Rock, AR; Proteomics Core Facility, UAMS, Little Rock, AR; Department of Nutrition, University of California, Davis, CA; Arkansas Children’s Nutrition Center and Department of Pediatrics, University of Arkansas for Medical Sciences, Little Rock, AR; Division of Nephrology, University of California, Irvine, CA; Division of Nephrology, UAMS, Little Rock, AR

## Abstract

**Background:** Resistant starch is a prebiotic metabolized by the gut bacteria. It has been shown to attenuate chronic kidney disease (CKD) progression in rats. Previous studies employed taxonomic analysis using 16S rRNA sequencing and untargeted metabolomics profiling. Here we expand these studies by metaproteomics, gaining new insight into the host-microbiome interaction.

**Methods:** Differences between cecum contents in CKD rats fed a diet containing resistant starch with those fed a diet containing digestible starch were examined by comparative metaproteomics analysis. Taxonomic information was obtained using unique protein sequences. Our methodology results in quantitative data covering both host and bacterial proteins.

**Results:** 5,834 proteins were quantified, with 947 proteins originating from the host organism. Taxonomic information derived from metaproteomics data surpassed previous 16S RNA analysis, and reached species resolutions for moderately abundant taxonomic groups. In particular, the *Ruminococcaceae* family becomes well resolved – with butyrate producers and amylolytic species such *as R. bromii* clearly visible and significantly higher while fibrolytic species such as *R. flavefaciens* are significantly lower with resistant starch feeding. The observed changes in protein patterns are consistent with fiber-associated improvement in CKD phenotype. Several known host CKD-associated proteins and biomarkers of impaired kidney function were significantly reduced with resistant starch supplementation. Data are available via ProteomeXchange with identifier PXD008845.

**Conclusions:** - Metaproteomics analysis of cecum contents of CKD rats with and without resistant starch supplementation reveals changes within gut microbiota at unprecedented resolution, providing both functional and taxonomic information. Proteins and organisms differentially abundant with RS supplementation point toward a shift from mucin degraders to butyrate producers.

## Introduction

Recent studies point to gut microbiome dysbiosis as one of the key contributors to the progression of chronic kidney disease (CKD) and its complications (1–3). During the course of CKD, gut dysbiosis increases and compromises the intestinal epithelial barrier, leading to leakage of microbial-derived toxins into the bloodstream and resulting in increased inflammation that may further exacerbate CKD (2). One suggested contributor to the dysbiosis is increased urea in intestinal fluids. Consequently, the urease-containing species proliferate in the gut, leading to damage of the epithelial barrier. Indeed, the CKD-associated microbiota have been characterized by an increase in bacterial species encoding for urease and uricase, and indole- and p-cresol producing enzymes, and depletion of microbes expressing short-chain fatty acid-forming enzymes (4).

Currently, CKD patients are often prescribed a diet that contains low quantities of fiber in order to limit the intake of potassium and avoid cardiac arrhythmias. However, in various models it has been shown that certain fibers can promote gut health and function, by increasing a gut microbiota population that dampens gut permeability and limits damage to the mucus layer caused by utilization of host glycans. Since CKD is a pro-inflammatory condition, and kidney damage may be exacerbated under conditions of gut microbiota dysbiosis, it is worth considering if increasing dietary fiber could help limit CKD complications and improve kidney function. One potential candidate to supplement a CKD diet is high-amylose maize-resistant starch type 2 (HAMRS2), a prebiotic which is metabolized by the gut microbes and has been shown to improve outcomes in a rat model of CKD (5,6). A previous study using taxonomic analysis characterized microbiome-related changes caused by resistant starch (RS) supplementation in CKD rats (5),. Microbiome and metabolomics data were correlated to identify potential metabolic pathways impacted by gut bacteria and linked to improvements in kidney function. In summary the previous study (5) provided strong evidence that resistant starch-induced microbiome shifts results in reduced inflammation and protection of the gut epithelial barrier.

Earlier studies in healthy humans, animals, and *in vitro* models showed increased levels of *Bifidobacterium, Ruminococcus, Lactobacillus, Bacteroides, Eubacterium, Allobaculum*, and *Prevotella* upon dietary supplementation of RS (7–11). Some of these organisms, e.g. *Ruminococcus bromii*, have been shown to contain genes for starch utilization and are proven direct degraders of RS (12–14). The organisms that increase in abundance upon RS supplementation are feeding on mono- and oligo- saccharides derived from RS-degradation by the direct degraders (12). *In vitro* stable-isotope probing followed by 16S rRNA sequencing of metabolically labeled RNA further validated that many of these bacteria utilize RS (15). Similar trends – showing increases in RS degraders and utilizers upon supplementation with RS in the rat model of CKD were observed by Kieffer et al. (5).

To gain further insight into the mechanism of RS action, we employed a metaproteomics analyses of the samples derived from the CKD rat model described previously (5,16). These analyses are capable of characterizing a complex protein mixture from an environmental sample. We employed three different quantitative proteomics techniques – absolute intensity-based quantification, spectral counting and TMT labeling, to characterize differences in metaproteome composition between RS-fed rats with CKD (CKD-RS), and the CKD rats fed with a host-digestible starch (CKD-DS).

## Methods

### Study design, animals and diets

We used the same animals as in (5). As indicated in that report, rats were randomized to receive semi-purified pelleted diets supplemented (59% by weight) with either the rapidly digestible starch (**DS**) amylopectin (low fiber) or HAMRS2 (Hi-Maize 260, Ingredion, Westchester, IL) (**RS**) for 3 wk (n = 9 rats/group). See **Supplemental Figure 1** and **Supplemental Text** for more information.

### Gel-LC MS, TMT-labeling, Basic HPLC, Tribrid-Fusion-Orbitrap Mass Spectrometry

Mass-spectrometry was performed by the UAMS Proteomics Core. Standard proteomics core protocols were adapted for this study (See **Supplemental Text** for detailed procedures and settings).

### De novo peptide sequencing

*De novo* sequencing was performed by PEAKS Studio v 8.0 (17), see **Supplemental Text** for detailed settings and parameters.

### Preliminary taxonomy analysis

*De novo* peptide sequences were submitted to the online metaproteomics tool UniPept (18,19) and to MetaCoMET (Metagenomics Core Microbiome Exploration Tool (20), to provide initial assessment of taxonomic diversity and sample quality (**Supplemental Text, Supplemental Figs. 2,3**).

### Data Sharing

The mass spectrometry proteomics data have been deposited to the ProteomeXchange Consortium via the PRIDE (21) partner repository with the dataset identifier PXD008845 and 10.6019/PXD008845

## Statistical Methods

For peptide identification and protein inference, a multi-step database search strategy was used in PEAKS Studio to arrive at the final list of identified proteins (see **Supplemental Text** for detailed description of the protein inference using the multi-step database strategy). For protein quantification, Scaffold v. 4 with quantitation module (Proteome Software) was used. Data and the custom fasta database were exported from PEAKS into Scaffold as mzIdentML and mascot generic format files. Scaffold-derived normalized spectrum abundance factors (NSAF) values and Total Unique spectral counts were used for the statistical analysis in R/bioconductor (22). Taxonomic information from the NCBI-nr database (downloaded Oct. 16, 2016) was used to derive taxonomic units. Sum of spectral counts matching a given taxonomic unit, weighted by a total spectral count per a given replicate, was used as a measure of abundance for that taxonomic unit. Moderated t-test (limma package, Bioconductor) was used to establish differential abundance of taxonomic units. Bonferroni-Hochberg correction for multiple testing was enforced both for protein and taxonomic unit quantification. See **Supplemental Text** for additional details

### Database search and Taxonomic analysis with MaxQuant and Andromeda and iBAQ quantification

In addition to spectral-count-based protein quantification in PEAKS, we performed parallel quantitative analysis with MaxQuant using both NSAF and iBAQ (Absolute Label-Free Protein Quantification) methods (see **Supplemental Methods** for detailed description). As a result, 2,842 proteins were quantified. Taxonomic analysis with spectral counts was performed in the same way as with PEAKS-derived proteins. Similarly, taxonomic units were quantified using intensity values by summing intensity values from iBAQ.

### Protein and taxonomic unit quantification using TMT labeling

PEAK Studio v. 8.0 with a quantitative module was used to analyze the TMT-labeled multiplexed experiments. 549 proteins and 149 taxonomic units were quantified using the TMT method. 45 taxonomic units were established as significant (p<0.05). See **Supplemental Text** for further details.

### Heatmaps Clustering, and PCA analysis

Heatmaps and hierarchical clustering of spectral count and TMT datasets were performed using the Bioconductor package ComplexHeatmap (23). PCA analysis was performed using base R.

## Results and Discussion

### Metaproteomics protein signature of cecal contents from rats fed Resistant Starch or Digestible Starch diets

Combining complementary quantification methods (TMT, PEAKS, and MaxQuant) 5,834 unique proteins were quantified in total (**Supplemental Table I, *tab (a*)**). The TMT method showed significant bias toward proteins and yielded significantly lower numbers of quantified proteins (452 vs. 3,007 quantified by PEAKS – **Supplemental Text**). The protein abundance signature clearly defines the phenotype induced by a resistant starch diet (**Figure 1**, *left panel*, **Supplemental Figure** 4). Notably, more host proteins are reduced in concentration upon resistant starch supplementation (*black lanes* indicate host proteins, while grey indicate bacterial proteins on the annotation bar next to the heatmap). This result is consistent with previous reports that indicate an increase in bacterial biomass upon resistant starch supplementation (3).

**Figure 1.**
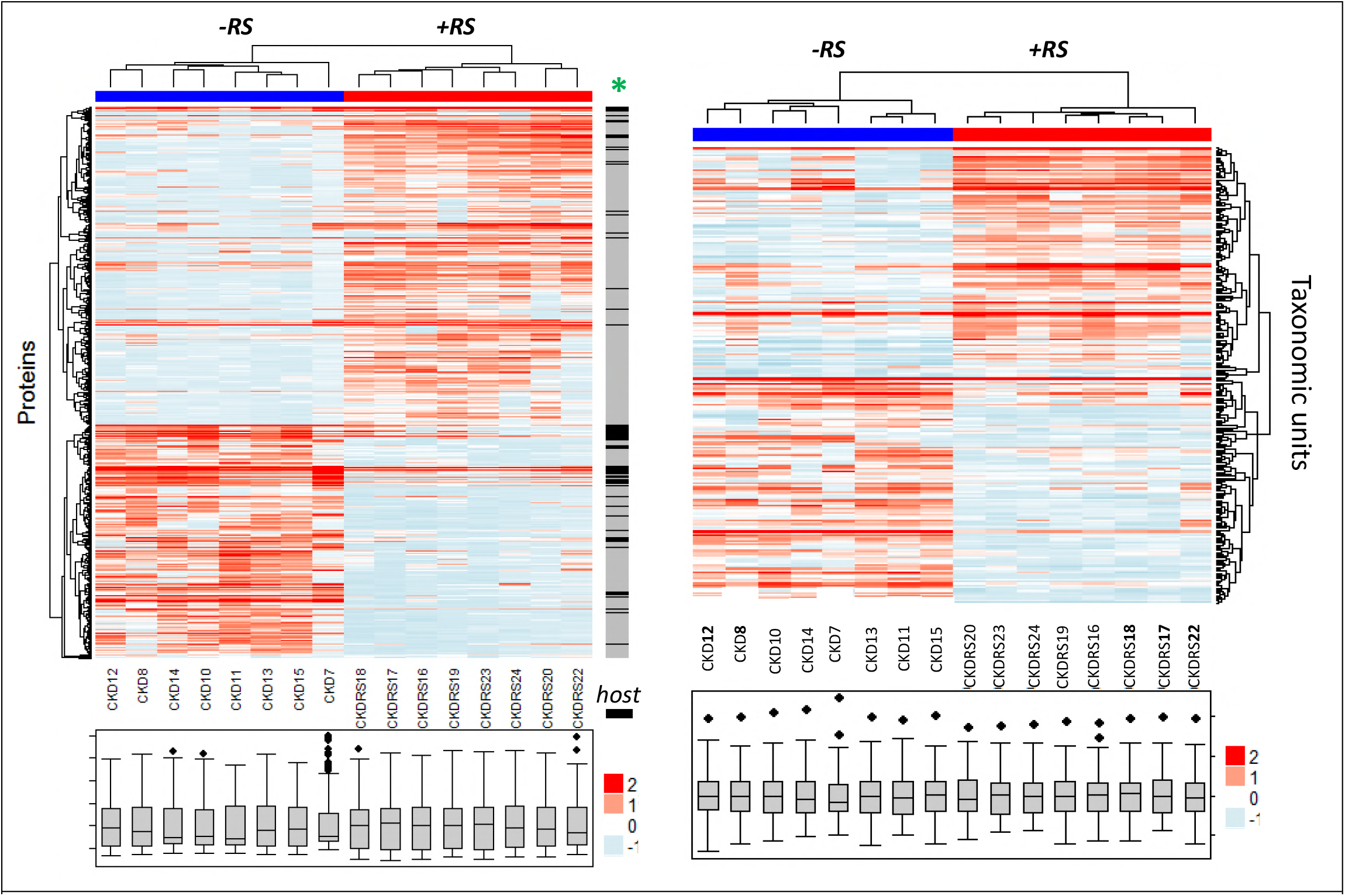
Phenotype induced by resistant starch in chronic kidney disease (CKD) rats is well-defined by metaproteomics data. *Left heatmap*, cecal content protein abundance values were averaged across three different labeling-free methods: PEAKS spectral counting, MaxQuant spectral counting and intensity based-quantification. The averaged values were log-transformed and normalized and plotted using ComplexHeatmaps R/Bioconductor package. 506 proteins, deemed significant by MaxQuant and by PEAKS are shown (p<0.01). CKD and CKDRS phenotypes are indicated by *blue* and *red* colors on the top heatmap annotation bar. Distribution properties are shown in BOX plots for each of the animals at the bottom, indicating similar distributions between different animals. The raw annotation bar is adjacent to the right edge of the heatmap – *“host”* indicates rat proteins by *black* color, and microbial proteins by *grey* color. *Right heatmap*, taxonomic units abundance values derived from the protein data were log-transformed and normalized and plotted using ComplexHeatmaps. Significant taxonomic groups (p <0.05) are shown.

Hierarchical clustering procedures applied to abundance values derived by metaproteomics separated the two phenotypes into two distinct clusters (**Figure 1**, *left heatmap*). This separation also held when all proteins were considered, not just differentially abundant ones (**Supplemental Figure 4**) We note that that if rats **9** (CKD group) and **21** (CKD-RS group), were included in the heatmap, it would break the clear separation of the two phenotypes and form a separate cluster (**Supplemental Figure 5**). Upon further scrutiny, these two samples showed significant degradation of bacterial proteins, and as a consequence, low-quality fragmentation mass spectra and high host-to-bacterial protein ratio (**Supplemental Figure 6**). These samples were therefore excluded and not used for the final analysis.

There were 179 host proteins that were differentially abundant between the cecal contents of CKD-DS and CKD-RS rats: 125 were proteins reduced in CKD-RS and 54 increased in CKD-RS (**Table I, Supplemental Table I** – complete protein list). Among the 125 host proteins lower in CKD RS-fed, approximately half were enzymes. The remaining proteins were related to humoral immune response, several proteins previously associated with epithelial–mesenchymal transition (e.g. thioredoxin, S100-A6) and several proteins, previously reported, directly or indirectly, to be associated with CKD (**Table I**, and see **Supplementary text** for more detail).

**Table I.** Representative differentially-abundant host proteins in cecal contents from CKD rats fed digestible starch (DS) or resistant starch (RS).

| **<u>Rat Proteins</u>** | **<u>Log2(CKD RS-fed: CKD)<sup>a</sup></u>** | **<u>p value<sup>b</sup></u>** | **<u>cited to be CKD-related</u>** |
| --- | --- | --- | --- |
| ***reduced*** |  |  |  |
| <u>thioredoxin</u> | -3.2, -1.7, -1.1, -1.5 | 2x10<sup>-3</sup> |  |
| <u>A1-microglobulin</u> | -3.9, nd, nd, nd, | 6x10<sup>-10</sup> | (24) |
| <u>haptoglobin</u> | nd, nd, -2.3, -2.7 | 5 x 10<sup>-3</sup> | (24, 25) |
| <u>calcineurin</u> | -3.0, -3.4, nd, nd | 1.8 x 10<sup>-5</sup> | (26) |
| <u>S100-A6</u> | Nd, -2.6, -3.7, -7.1 |  |  |
| <u>calreticulin</u> | -1.6, -3.9, -3.6, -6.1 | 3x10<sup>-5</sup> | (27) |
| <u>Angiotensin-converting enzyme</u> | nd, nd, -2.5, -4.3 | 1x10<sup>-3</sup> | Well-established therapeutic target in CKD(28) |

| **increased** |  |  |  |
| --- | --- | --- | --- |
| Annexins A7, VII,A2 | 2 to 3 <sup>c</sup> | 10<sup>-3</sup> to 10<sup>-8</sup> <sup>c</sup> | (29) |
| voltage-dependent anion-selective channel protein 2 | 1.6, 4.5, 2.8, 5.2 | 2x10<sup>-6</sup> | (30, 31) |
| Claudin-3 | nd, nd, 2.0, 3.2 | 9x10<sup>-3</sup> |  |

Among the 54 host proteins that were increased in the CKD-RS group, about one third were enzymes. The largest group of proteins with similar functions was related to the immunoglobulin families. Another group included annexins (**Table I, Supplementary Table I**) that function as intracellular Ca^2+^ sensors and participate in cellular membrane repair (29), suggesting that in CKD-RS rats a process of intestinal cell repair, presumably damaged by bacteria and/or bacterial metabolites, is ongoing. Other small groups of proteins with similar function included a group of voltage-dependent ion channel proteins that regulate fluxes across the outer mitochondrial membrane and sodium pump subunit proteins. Mitochondrial impairment has been shown in CKD patients and animal models, particularly in the form of a decrease in mitochondrial DNA and down-regulation of many mitochondrial genes and proteins (30,31). Down-regulation of voltage channel proteins in CKD rats could be, in part, causing the increase of oxidative stress in CKD. Similar to previous studies (3,16), we observed an increase in the tight junction protein claudin-3 (2.6 fold change in CKD-RS).

Interestingly, many of the individual proteins we found to be differentially abundant between CKD-DS and CKD-RS rats were previously identified in other reports on CKD and kidney associated diseases (16,24,26,27,31,32). Importantly, these proteins were identified either in plasma or in urine. Identification of these proteins in cecal contents points to a potential utility of stool samples as a source of CKD protein biomarkers; however, the pathophysiological link between the association of the proteins in the GI tract with resistant starch and their association in urine and plasma during CKD is currently unclear.

### Molecular processes influenced by dietary resistant starch supplementation in the cecal contents of CKD rats

To define enrichment of functional patterns associated with the differentially abundant proteins found by metaproteomics, we performed Gene Ontology analysis using Blast2Go (33). The 179 differentially-abundant host proteins described above were combined with 1198 differentially abundant bacterial proteins, the latter of which included 359 reduced and 839 higher proteins with RS feeding. The list of proteins differentially abundant between CKD-DS and CKD-RS rats was tested for enrichment of GO terms (biological process, molecular function and cellular compartment) for bacterial and host proteins separately using Fisher exact test. All GO terms significant at the FDR level of 0.05 were considered significantly enriched.

Biological processes under-represented in the CKD-RS list of rat proteins were related to known biological processes occurring in CKD, namely aldehyde metabolism (indicative of lipid peroxidation processes), and humoral immune response. GO categories significantly over-represented in CKD-RS rat proteins were related to the sodium-potassium pump subunits and cross-membrane transport; in theory this may indicate that control of pH, osmotic pressure and cell volume are compromised in CKD rats and improve with the RS-rich diet. However, the mechanisms of sodium-potassium pump regulation are complicated: for example, it was suggested that cardiotonic steroids mediate signal transduction through the Na/K-ATPase, and its downregulation could be indirectly implicated in profibrotic pathways (34). Given that it is difficult to predict the consequences of insufficient sodium pump levels without further experiments, this observation warrants future study. For other GO analysis details see **Supplemental Text**.

Many biological processes and molecular functions, associated with CKD, especially at the more advanced stages, were previously characterized by transcriptomics. These included down-regulation in the kidney of regulatory proteins involved in cytoskeleton organization, microtubule assembly and stability, epithelial–mesenchymal transition, extracellular matrix remodeling, cell motility and migration, cell adhesion, apoptosis, cell differentiation, proteolysis, aminoglycan metabolic process and protein N-linked glycosylation (35,36). One advantage metaproteomics offers is the simultaneous analysis of bacterial and host proteins, with metaproteomics offering higher resolution of bacterial portion of the proteome compared to metatranscriptomics. Indeed, in its current stage dual transcriptomics (i.e. simultaneous analysis of host and microbiome transcripts) is more challenging than metaproteomics. The pipeline for dual RNA-seq analysis has the same building blocks as conventional RNA-seq pipeline: reads must be cleaned, mapped, normalized and differentially expressed transcripts identified. However, as recommended in (37) host and pathogene reads should be mapped to the reference genomes and analyzed separately (37). The latter creates a difficulty of using additional READemption pipeline for mapping bacterial reads, in addition to conventional TopHat pipeline for mapping host reads. READemtion pipeline uses the short read mapper segemehl and its remapper lack (38,39) unlike Tophat that uses bowtie (40). Also, when ones choses between transcriptomics and proteomics to characterize a disease – in our opinion proteomics is a better choice because 1) cellular phenotypes are defined by proteins more so than by transcripts, hence proteins are better biomarkers 2) druggable targets are usually proteins and not transcripts 3) correlation between protein abundance and transcript abundance is typically 40% to 60% depending on the cell type.

For bacterial proteins, GO categories over-represented in CKD-RS samples (biological processes, molecular functions and cellular compartments) were all related to active bacterial proliferation, emphasizing a shift in CKD-RS microbiota toward new actively dividing bacterial populations, which are able to thrive on RS (see **Supplementary Text** for more detail). Active bacterial proliferation can also partially explain the previously observed reduction in harmful tryptophan and tyrosine metabolites in CKD-RS rats (indoxyl and p-cresol, respectively). Instead of conversion to the toxins, tryptophan and tyrosine could be incorporated into newly synthesized proteins.

The analysis of significantly under-represented GO categories in CKD-RS for bacterial proteins supports the idea that in CKD-DS samples there is an ongoing process of mucin degradation that is more active than in CKD-RS samples. It could indicate preferential foraging on mucin proteins by gut microbiota in CKD-DS rats when compared to CKD-RS rats. In our list of bacterial species, under-represented in CKD-RS, several mucin degraders were evident (e.g. *Mucispirillum schaedleri, R.gnavus, R.torques* – **Supplemental Table II**), confirming the idea that bacteria reduced in CKD-RS are preferentially mucin degraders, or generalists. For example, *R.torques* which is increased in CKD-RS can utilize both mucin and amylose as a major source of carbohydrates. Thus, the taxonomic units inferred from metaproteomics data clearly define the resistant starch phenotype. **Supplemental Table II** summarizes the taxonomic information inferred from metaproteomics experiments. Hierarchical clustering of the taxonomic unit abundance data separates the two phenotypes (**Figure 1**, *right*), similar to the protein-level data.

### Cecal microbiome composition revealed by metaproteomics at high resolution

#### Alpha-diversity

Using metaproteomics data for the inference of taxonomic units, we observed increases in alpha diversity for the samples derived from the CKD-RS rats, when the lowest level of taxonomy is considered. iBAQ and SAF method of quantification gives a 50% (p=0.013) and 30% increase (p=0.008), respectively (**Supplementary Text, Supplementary Figure 7**). This increase in alpha diversity is opposite to what has been reported in previous studies of resistant starch supplementation in healthy pigs where 16S RNA was used for the taxonomic inference (11); it is also opposite to what has been inferred from the same rats in Kieffer et al. using 16S RNA (5). The reason for this discrepancy has to do with taxonomic resolution: the diversity drops when only family and genus taxonomic levels are considered (limit of resolution for 16S RNA method), but the diversity within specific families increases at the species level (see example for <u>*Ruminococcus* genus</u> below). Thus, our data using metaproteomics support an increase in microbial diversity in response to RS diet.

#### Changes in gut microbial composition

As with the 16S RNA studies, we observe an increase in the *Bacteroidetes-to-Firmicutes* ratio in CKD-RS group. However, because of the species-level resolution the absolute numbers differ. In the previous analysis using 16S RNA, an overall ~2.5 times increase in *Ruminococcus genus* was observed. Metaproteomics analysis provides species-level resolutions and as a result, some species of *Ruminococcus* dramatically increase and some dramatically decrease with the RS supplementation. For example, *Ruminococcus bromii* is the key degrader of resistant starch in the mammalian gut (12). According to the metaproteomics analysis (Supplemental **Table I**, *tab (c*), **Supplemental Table II**), *Ruminococcus bromii L2-63* is increased ~12 times upon addition of resistant starch. At the same time, some species of this genus were decreased. For example, *Ruminococcus albus* decreased 20 fold and *Ruminococcus flavefaciens* decreased 10 fold. *R. albus, R.gnavus* and *R.flavefaciens* are fibrolytic bacteria that are able to process complex plant polysaccharides by their cellulolytic and hemicellulolytic enzymes. They use ammonia almost exclusively as their source of nitrogen. These bacteria are also known to thrive under high pH (above 6.0). Concordantly, in the preceding study pH was shown to drop from 8 to 6.75 in the CKD-RS rats (5,16). In contrast, *R. bromii* and *R.torques* are amylolytic and use RS as the main source of nutrients. Amylolytic (resistant starch degraders) and fibrolytic (cellulose degraders) species split the *Ruminoccocus* genus into two groups. Strikingly, the taxonomic resolution of metaproteomics analysis allowed us to observe clearly that fibrolytic *Ruminoccocus* are reduced with dietary RS, while amylolytic species are increased with the addition of RS. Beyond the biological importance of this observation per se, it demonstrates the power of metaproteomics analysis and the lower resolution of the 16S RNA method.

#### Potential mechanisms of RS action

The question that remains to be fully understood is the mechanism(s) through which the RS diet exerts such a dramatic shift in control of biological processes and functions that are different between CKD-DS and CKD-RS rats. In part, it might be explained by bacterial population that shifts from mucin foraging to RS as a source of nutrients, relieving the host system from the constant flow of toxins that traverse the gut barrier. Other effects may involve reduction of oxidative stress. In the past, it was shown that kidneys from CKD patients have an impaired mitochondrial respiratory system (31), decreased DNA mitochondrial copy number (30), loss of mitochondrial membrane potential and lower ATP production (41). The hypothesis that new bacterial populations influence mitochondrial biogenesis and activity (for example, through the influx of short chain fatty acids) is an attractive one. Recently it was shown that butyrate supplementation improved mitochondrial biogenesis in mice (42), presumably via inhibition of histone deacetylase (HDAC) that may down-regulate expression of PGC-1α, associated with mitochondrial dysfunction (43–45). There is growing evidence that RS prevents colonic DNA damage via the production of SCFA, especially butyrate (46,47). In fact, it has been known, for almost a decade, that butyrate has a central role in maintaining gut epithelial integrity via involvement in key biological processes, such as being a source of energy for colonocytes, promoting fatty acid oxidation, having anti-inflammatory activity, limiting oxidative stress and inducing cell cycle arrest (48,49). In all these processes butyrate putatively functions by blocking substrate access to active sites in HDACs. Butyrate is a microbial fermentation product and butyrate producers are polyphyletic, belonging to different bacterial species, most known are members of *Lachnospiraceae* and *Ruminococcaeae* (50). Butyrate is synthesized by those microorganisms via pyruvate and acetyl-coenzyme A (CoA) by breakdown of complex polysaccharides (such as RS). We observed that RS supplementation, at least for the *Ruminococcaeae* family, results in a shift from fibrolytic family members to amylolytic family members that are well known butyrate-producers (*Ruminococcus bromii, Ruminococcus torques).* Typically, in the absence of sufficient dietary fibers, commensal and pathogenic bacteria start to forage on mucin glycans to harness carbon and energy (51,52). While we did not examine if microbes lowered with RS feeding are mucin degraders we did find indirect evidence in support of that from GO categories reduced with RS feeding (monosaccharide metabolic process and L-fucose catabolic process). We therefore hypothesize that there is a relative shortage of fiber in CKD-DS and the RS diet shifts gut microbial communities, from bacteria foraging on mucins and ammonia and contributing to leaky gut phenotype to bacteria utilizing RS instead, and producing butyrate as a byproduct of fermentation.

Recently it was found that out of 3,184 sequenced bacterial genomes, mostly from the Human Microbiome Project, 225 were likely to be butyrate producers (50). From the list of bacteria, up-regulated in CKD-RS, only *Eubacterium rectale, Clostridium botulinum and Lachnospiraceae bacterium* strains are present in this list of potential butyrate producers, presumably because rat and human gut microbiomes differ. It would be interesting to evaluate quantitatively the amount of mucin degraders and butyrate producers caused by the diet shift by genomics and transcriptomics, to reduce biases due to proteomics undersampling (e.g. bias against low abundance proteins).

To summarize, dietary RS supplementation in rats ameliorates chronic kidney disease coincident with a massive shift in gut microbial communities (**Figure 2**). Identified organisms and proteins point toward a higher population of butyrate-producing bacteria, and reduced abundance of mucin-degrading bacteria. It is speculated that the bacterially-derived butyrate leads to improvement of oxidative stress and inflammation, as well as to improvement in other biological processes otherwise impaired in CKD. In addition, it is hypothesized that gut barrier function through maintenance of the mucin barrier may also play a role in RS-associated improvements in CKD phenotype. Finally, resistant starch supplementation leads to the active bacterial proliferation and the reduction of harmful bacterial metabolites. The fact that simple change of available source of nutrients (from ammonia/mucins to RS) leads to system-level changes underscores the importance of diet during disease management and highlights the potential role of the gut microbiome in disease progression.

**Figure 2.**
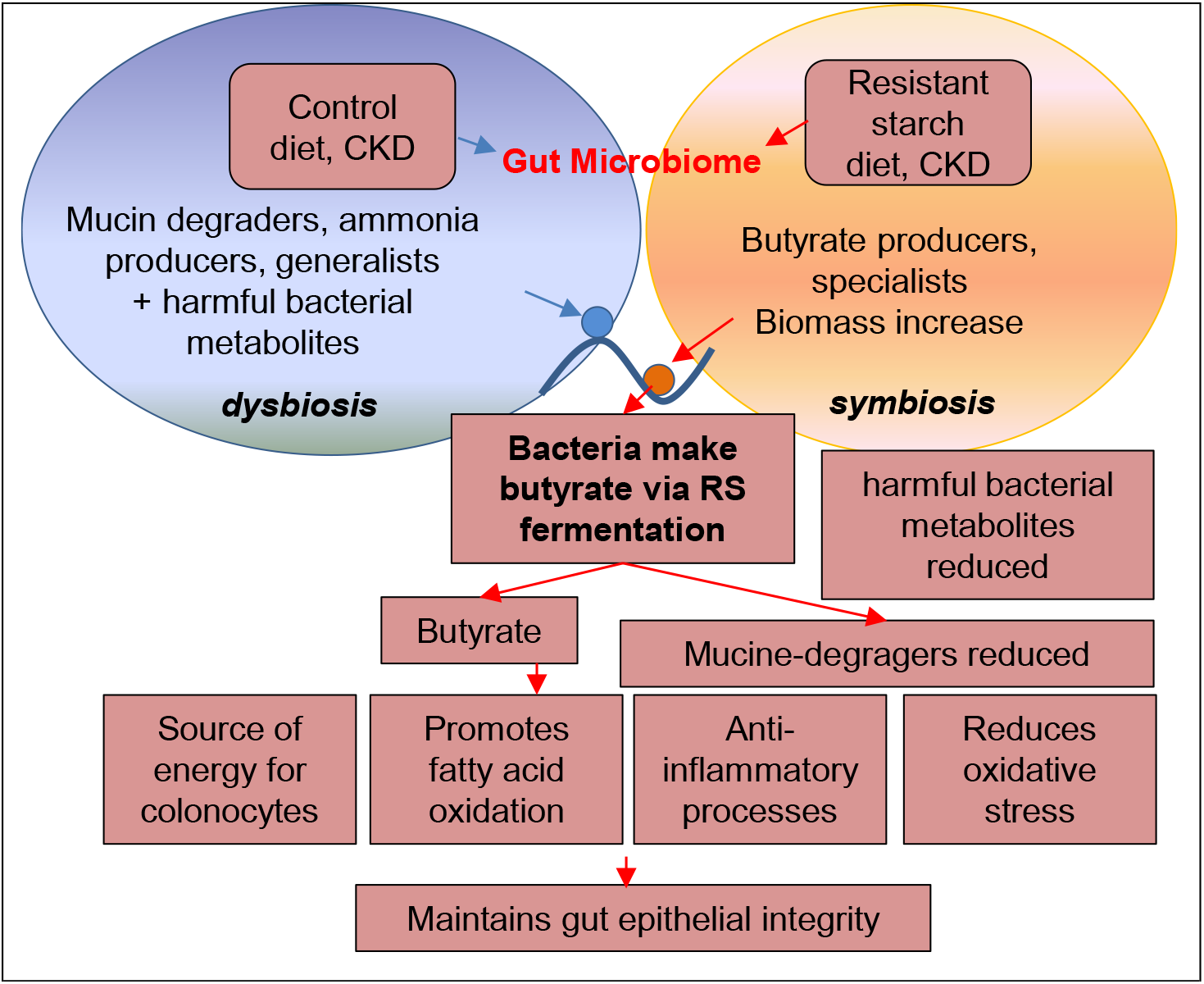
Working model of how RS diet changes gut microbiome composition, and in turn alleviates CKD.

